# Mobile Element Insertions and Associated Structural Variants in Longitudinal Breast Cancer Samples

**DOI:** 10.1101/2020.02.10.942680

**Authors:** Cody J. Steely, Kristi L. Russell, Julie E. Feusier, Yi Qiao, Sean V. Tavtigian, Gabor Marth, Lynn B. Jorde

## Abstract

While mobile elements are largely inactive in healthy somatic tissues, increased activity has been found in cancer tissues, with significant variation among different cancer types. In addition to insertion events, mobile elements have also been found to mediate many structural variation events in the genome. Here, to better understand the timing and impact of mobile element insertions and structural variants involving existing mobile elements in cancer, we examined their activity in longitudinal samples of four metastatic breast cancer patients. With whole-genome sequencing data from multiple timepoints through tumor progression, we used mobile element detection software followed by visual confirmation of the insertions. We identified 11 mobile element insertions or structural variants involving existing elements and found that the majority of these occurred early in tumor progression. Two of the identified insertions were SVA elements, which have rarely been found in previous cancer studies. Most of the variants appear to impact intergenic regions; however, we identified a translocation interrupting *MAP2K4* involving *Alu* elements and a deletion in *YTHDF2* involving mobile elements that likely inactivate reported tumor suppressor genes. The high variant allele fraction of the *MAP2K4* translocation, the loss of the other copy of *MAP2K4*, the recurrent loss-of-function mutations found in this gene in other human cancers, and the important function of *MAP2K4* indicate that this translocation is potentially a driver mutation. Overall, using a unique longitudinal dataset, we find that most variants are likely passenger mutations in the four patients we examined, but some variants impact tumor progression.

## Introduction

Mobile elements, or transposable elements, are segments of DNA that are capable of mobilizing from one genomic location to another. These elements compose a significant portion of the human genome, with estimates ranging from nearly 50% to 66% (Lander et al. 2001; de Koning et al. 2011). In humans, retrotransposons represent a class of active mobile elements, inserting new elements through a “copy and paste” mechanism (Boeke et al. 1985). While most of these elements have become transcriptionally inactive over time (Brouha et al. 2003; Beck et al. 2010), three mobile element families remain active in humans, including LINE-1 (L1), *Alu* elements, and SVA. These three mobile element families compose nearly one-third of the human genome (Lander et al. 2001; Xing et al. 2013). With only a small fraction of these elements retaining the ability to retrotranspose, recent work has shown that germline mobile element insertions in humans are quite rare (Feusier et al. 2019). The majority of these insertions seem to have no adverse impact; however, some insertions have been found to cause disease (Hancks and Kazazian 2016). Mobile element activity in human disease became a subject of interest after two hemophilia A patients were found to have *de novo* L1 insertions in the *F8* gene (Kazazian et al. 1988). Since this initial discovery, mobile element insertions have been found to be associated with more than 130 disease cases (Kazazian and Moran 2017).

Somatic mobile element insertions have not been identified in many healthy tissues, though detection of these low-frequency events is difficult without sufficient coverage. An exception to this is in neurons, where somatic mosaicism of L1 insertions have been identified (Muotri et al. 2005; Coufal et al. 2009; Richardson et al. 2014; reviewed in Erwin et al. 2014). This increase in activity may be due to slight changes in methylation at L1 loci (Coufal et al. 2009). Recently, multiple studies have noted varying degrees of activity of mobile elements in numerous cancer tissues (Lee et al. 2012; Helman et al. 2014; Tubio et al. 2014; Doucet et al. 2015; Doucet-O’Hare et al. 2016; reviewed in Burns 2017; Li et al. 2020; Rodriguez-Martin et al. 2020), with some analyses including metastatic samples (Rodić et al. 2014; Tubio et al. 2014; Ewing et al. 2015). The level of activity has been found to be quite variable in patients with the same type of cancer and also shows a high degree of variability among cancer types (Helman et al. 2014; Tubio et al. 2014). Regardless of cancer type, roughly half of all tumors have at least one somatic L1 insertion. Breast cancer patients with L1 activity were found to often have a single L1 insertion, with a small number of patients showing up to five insertions (Tubio et al. 2014).

The impact of mobile elements on the cancer genome is not limited to somatic insertions but also includes the structural variation (SV) events associated with existing mobile elements (reviewed in Cordaux and Batzer 2009). Notably, mobile elements mediate roughly 10% of all SV events larger than 100 base pairs in the human genome (Xing et al. 2009). Mobile elements have been found to mediate SV events that can lead to cancer development (Mauillon et al. 1996; Hsieh et al. 2005). For example, approximately 42% of the *BRCA1* gene is composed of *Alu* elements (Welcsh and King 2001), making this gene a target for non-allelic homologous recombination, specifically *Alu-Alu* associated events (Petrij-Bosch et al. 1997; Puget et al. 1997; Rohlfs et al. 2000; Peixoto et al. 2013). Additional mobile element events have been classified in breast cancer as well (Morse et al. 1988; Walsh et al. 2006). L1 transduction events have also been examined in a variety of cancers (Tubio et al. 2014).

Multiple studies have shown that L1s, in particular, demonstrate increased retrotransposition activity in cancer tissues. However, the timing of these insertions or mobile element associated SVs has not been thoroughly investigated. Additionally, L1s and the structural variants that they mediate have been the focus of most previous studies, though *Alu* and SVA insertions or the SVs they mediate have been targeted in some studies (Lee et al. 2012; Tubio et al 2014; Rodriguez-Martin et al. 2020). In this study, we utilize longitudinally sampled breast cancer tissues to better understand the timing and significance of increased mobile element activity on the cancer genome.

## Results

To analyze mobile element activity during tumor progression, we used longitudinal whole-genome sequencing (WGS) data from four metastatic breast cancer patients. Tumor samples were obtained from these patients over 2-15 years with varying WGS and bulk RNA-seq timepoints (2-6) available for our analyses. Many of the tumor samples were collected from ascites and the surrounding pleural fluid. Tumor cells in ascites are descended from the primary tumor, though they may contain additional mutations and are likely to be polyclonal. Using ascites to analyze tumors may lead to improved characterization of the tumor and of the clonal changes that occur (Choi et al. 2017; Husain et al. 2017). Germline DNA was sequenced from the blood of each patient. The WGS timepoints for each patient, and a summary of treatment information over time, are shown in Supplemental Figure S1. More detailed treatment information for each patient can be found in the supplementary information of Brady *et al*. 2017. All four patients were estrogen receptor (ER) +. Patients 1 and 2 were human epidermal growth factor receptor 2 (HER2) +, while Patients 3 and 4 were HER2-.

We analyzed each of the four patients for mobile element insertions utilizing three tools (MELT, TranSurVeyor, and RUFUS, further described in the Methods section). After identifying and filtering mobile element insertions and SVs (see Methods), we generated a list of potential insertion sites. These potential insertion sites were compared to the matched germline sample for each patient to ensure that the identified insertion was a somatic event. Timepoints for each patient were analyzed individually and compared to the matched germline sample. Following our identification and filtering steps, we were able to identify mobile elements and associated variants at both very high Variant Allele Fraction (VAF) (100%) and very low VAF (∼5%). While the three tools used in this study generally identified the same insertion events, MELT and RUFUS excelled at identifying *Alu* elements, and TranSurVeyor found an additional L1 insertion that was not identified by the other tools. Our detection methods have likely not produced a comprehensive list of all SVs involving mobile elements due to the difficulty in detecting these variants in short-read sequencing data.

Collectively, we identified seven mobile element insertions (either classical or non-classical) and four structural variants involving mobile elements (Table 1). These variants all appear to be somatic mutations acquired during tumorigenesis or during tumor progression because these variants are not present in the corresponding germline samples. IGV images with schematics are shown for three of the events that we identified in Figure 1, with the remaining IGV images shown in Supplemental Figure S2 (schematics of the remaining events are included as Supplemental Figures S3, S4, and S5). We find that the majority of insertions and variants associated with mobile elements (nine of the eleven) occur before the first sampling timepoint because these were already present in the first and then all subsequent samples. The two variants that were not present in the first sampling timepoint were both SVA insertions. Both SVA insertions identified were present at low frequency in these patients. We also find high variability in the number of insertions and variants that were identified in each of these patients, with three of the patients (Patients 1, 2, and 4) showing multiple insertions and variants and Patient 3 showing only a single structural variant associated with mobile elements.

**Table 1.**
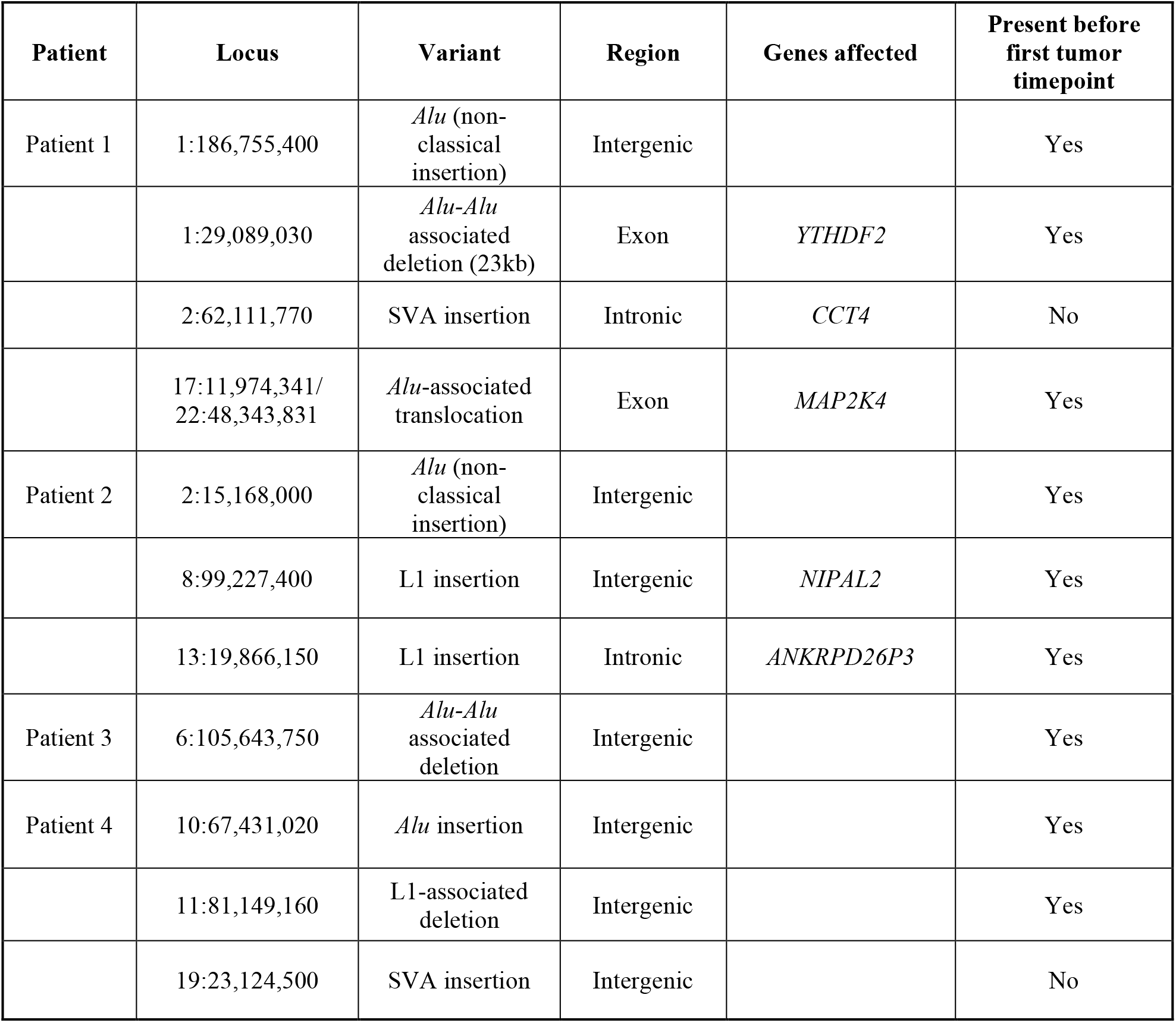
Mobile element insertions and structural variants involving mobile elements identified in four longitudinal breast cancer patients. Insertion or variant sites found within genes (exonic or intronic) are indicated in the “Genes affected” column.

**Figure 1.**
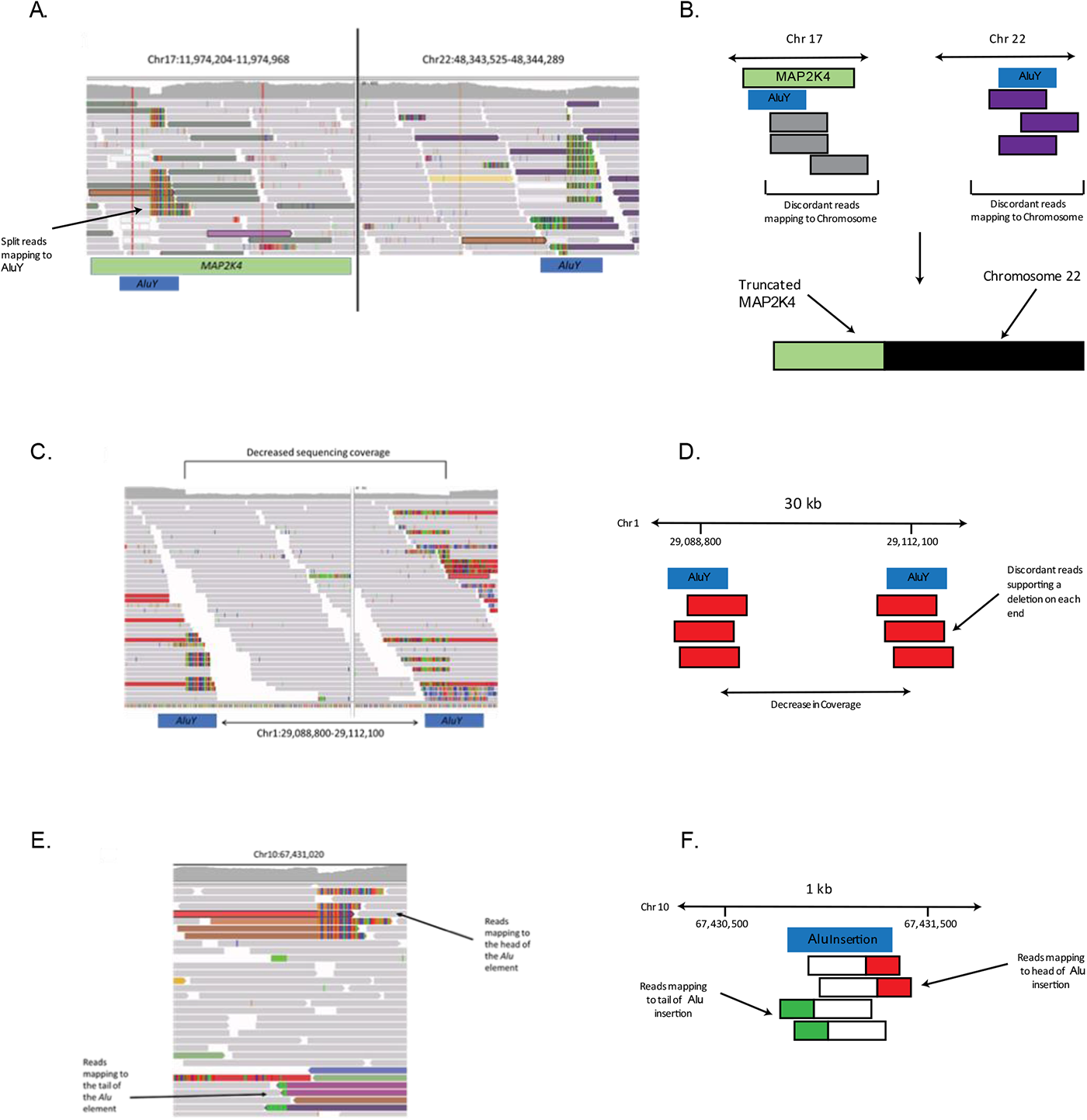
IGV images and schematics of some of the identified insertions and structural variants.A) Image of the translocation between Chromosomes 17 and 22 in Patient 1 that interrupts *MAP2K4*. Discordant reads on each chromosome map to the other end of the translocation. The split reads on Chromosome 17 map to a non-reference *Alu* element. The *Alu* element suspected to be involved in the translocation on Chromosome 17 (from the split reads) is shown as a blue box, and the reference *Alu* involved with the translocation on Chromosome 22 is also shown as a blue box. B) A schematic showing the translocation event (not to scale). The *Alu* elements associated with the translocation are shown as blue boxes. The discordant reads are shown as gray or purple boxes. A schematic of the translocation is shown below the arrow. C) Deletion involving *Alu* elements in Patient 1. *Alu* elements involved in the deletion are shown as blue boxes below the image. D) A schematic of the deletion involving two *Alu* elements. Each *Alu* element is shown in blue, with the discordant reads shown in red. E) Somatic *Alu* element insertion in Patient 4. The reads that map to the head and tail of the element are labeled by arrows. F) A schematic of the *Alu* insertion on Chromosome 10. The approximate location of the insertion (blue box) is shown with the split and discordant reads mapping to the head of the *Alu* shown in white and red boxes, while the split and discordant reads mapping to the tail of the *Alu* are shown in white and green boxes.

The use of multiple tools and visual inspection of the identified insertions excluded a large number of false positive calls. In total, MELT identified 8,825 potential mobile element variants, and TranSurVeyor identified 69,394 potential mobile element variants. For MELT, we analyzed every polymorphic variant (regardless of genotype) to determine if it was unique to the tumor sample. For TranSurVeyor, we did not use the built-in filtering mechanism to ensure that low frequency variants would be included for further examination. Following this filtering, we retained only those variants that passed visual inspection (See Methods). These 11 variants are shown in Table 1. Counts for the images produced by RUFUS were not included here as RUFUS is designed to detect many types of variants, not exclusively mobile elements.

Most of the identified insertions and variants were found in intergenic regions and are unlikely to impact gene expression as they do not overlap known regulatory or ultra-conserved regions. However, we identified multiple insertions that appeared to impact intronic regions: an SVA insertion in *CCT4*, which was identified in Patient 1, an L1 insertion in *NIPAL2* in Patient 2, and an L1 insertion in *ANKRPD26P*, which we identified in Patient 2. We also identified two *Alu-*associated somatic structural variation events that impact exons in two genes: an *Alu-Alu* associated deletion in *YTHDF2* and a translocation involving *Alu* elements that impacted *MAP2K4*. Both of these *Alu*-associated structural variants occurred in Patient 1 and occurred before the first sampling timepoint but were not present in the germline sequencing data.

To remove contamination from non-tumor cells, we estimated the tumor purity (or cancer cell fraction) of each timepoint. The purity estimates, as well as estimates for genome-wide copy number for each timepoint in each patient, are included in Supplemental Table S1. There are many changes in genomic copy number, with genome-wide estimates ranging from 1.86, just below normal cellular copy number, up to 4.12, more than twice that seen in most normal cells.

In these patients, tumor purity ranged from 30.91% to 94.5%, with a mean value of 76.25%. The only formalin-fixed, paraffin-embedded (FFPE) sample in the dataset had the lowest tumor purity value.

We calculated the VAF for each of the mobile element insertions or SVs identified after adjusting for tumor purity (Figure 2). The VAFs calculated from the adjusted data are slightly higher, but very similar to the VAFs calculated from the unadjusted VAFs (Supplemental Figure S6). Of the four insertions and SVs identified in Patient 1 (Figure 2A), the deletion in *YTHDF2* involving *Alu* elements, and the translocation in *MAP2K4* were present at very high frequency (>80% for the deletion; 100% for the translocation) at the first timepoint, with decreased frequency over time. The other two identified events were insertions (an *Alu* and SVA) and were present at much lower frequencies. The SVA identified in this patient was not present at the first two sampling timepoints but was present by the third and remained in the tumor cells in the fourth sampling timepoint. Each of these events show a slight decrease over sampling timepoints. We identified three somatic insertions in Patient 2 (Figure 2B), two L1 insertions, and one *Alu* insertion. The L1 insertion on Chromosome 13 was present at a higher frequency than the other two insertions found in this patient. The frequency of these variants closely resembles the frequency of the mobile element insertions seen in Patient 1, but does not reach the high frequency seen in the translocation or deletion of Patient 1.

**Figure 2.**
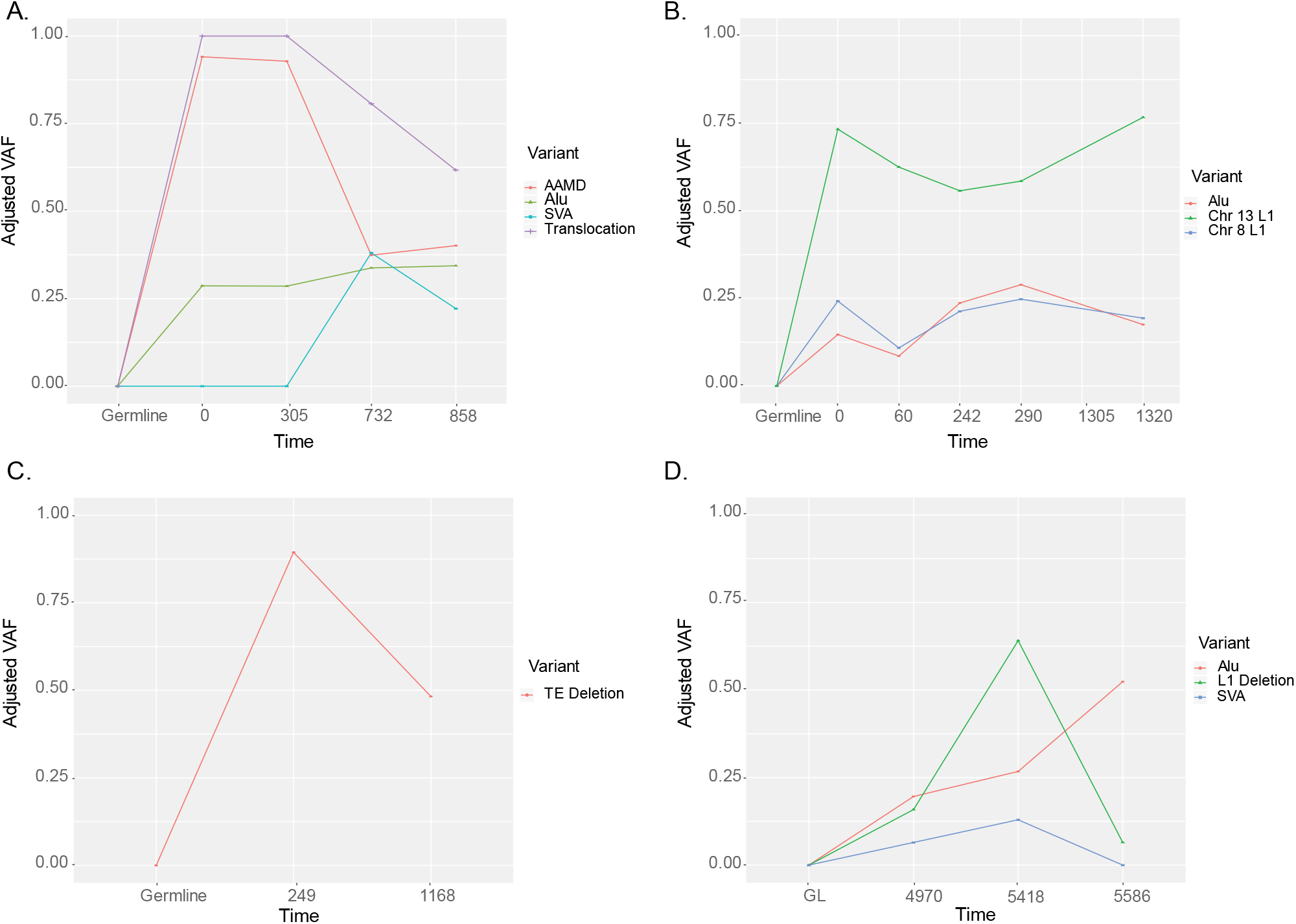
Adjusted VAF for each of the 4 patients. Parts A-D correspond to patients 1-4, respectively. The VAF for each patient has been adjusted to better reflect the percentage of cells in the sample that appear to be from the tumor. VAF is shown on the Y axis with each sampling timepoint for a particular patient shown on the X axis.

Of the four individuals sampled, Patient 3 (Figure 2C) had the fewest variants, a single deletion involving *Alu* elements. This deletion was present at a high frequency in the first sampling timepoint (about 75% of reads), but decreased to approximately 50% by the second sampling timepoint. In Patient 4 (Figure 2D), we identified two insertions, an *Alu* and SVA, and a single mobile element associated SV, a L1-L1 associated deletion. The first sequenced sampling timepoint for Patient 4 has been excluded in the VAF plot as it was a FFPE sample, decreasing our ability to accurately calculate a VAF for this timepoint. The SVA insertion identified in this patient was not present for the first sampling timepoint but was present for the next two sampling timepoints. By the final sampling timepoint, the SVA was no longer observable in the sequencing data. The L1-L1 associated deletion trends upward in VAF before sharply falling at the final timepoint. The *Alu* insertion in this patient maintains a steady VAF of ∼20%.

After identifying the mobile element insertions and the SVs that they mediate in these four patients (Table 1), we analyzed the impact of these variants on the cancer genome. Because both variants that impacted exons were found at high frequency in Patient 1, we examined the genomic copy number present in these regions. Specific genome-wide copy number estimates for the first timepoint in the genome of Patient 1 are shown in Figure 3. Both the deletion involving *Alu* elements and the *Alu* associated translocation appear to have been reduced to only a single copy, contributing to the high frequency for both.

**Figure 3.**
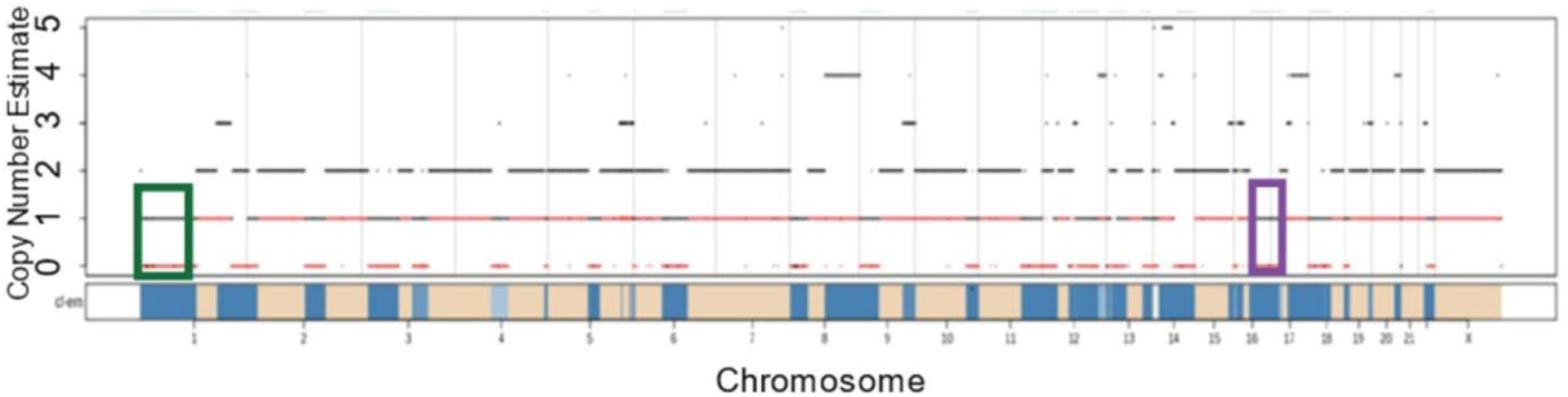
Copy number estimates from FACETS for Patient 1. A decrease in copy number along part of Chromosome 1 is shown in the green box. This green box includes *YTHDF2*, a gene that is partially deleted by a mobile element associated event. The purple box is highlighting a decrease in copy number along Chromosome 17. Included in this purple box is *MAP2K4*, a gene that is interrupted by a translocation event associated with mobile elements. The black lines in the figure represent the total copy number, while the red lines show the minor copy number for each segment.

Due to decreased copy number at the *MAP2K4* locus, and the translocation interrupting the one remaining copy of *MAP2K4* (shown in Figure 4A), we validated absence of the complete transcript and protein in the cancer cells of Patient 1. Creating a *de novo* transcript assembly of RNA-seq data for each timepoint in Patient 1 (see Methods), we find only truncated and hybrid transcripts of *MAP2K4* (Figure 4A; Sequences for these transcripts are shown in Supplemental Table S2. To further validate a lack of protein production by *MAP2K4* in the tumor cells of Patient 1, we performed a Western blot using MCF7 cells as a control and tumor cells taken from an ascites sample from Patient 1 (Figure 4B). While the control cells show clear production of MAP2K4, there appears to be a decrease in MAP2K4 in the tumor cells of Patient 1. The remaining production of MAP2K4 may be due to non-tumor cells in the sample. Further, mapping the transcripts from the RNA-seq data in Patient 1 (see Methods), we find an average of only 2.92 TPM that map to *MAP2K4*. The number of transcripts aligning to *MAP2K4* in Patient 1 is much lower than the median value found by GTEx (median 18.05 TPM; the GTEx Portal on 08/28/2020). Analyzing RNA-seq data from another patient that was not suspected to have any disruption to *MAP2K4*, we found an average of 14.80 TPM.

**Figure 4.**
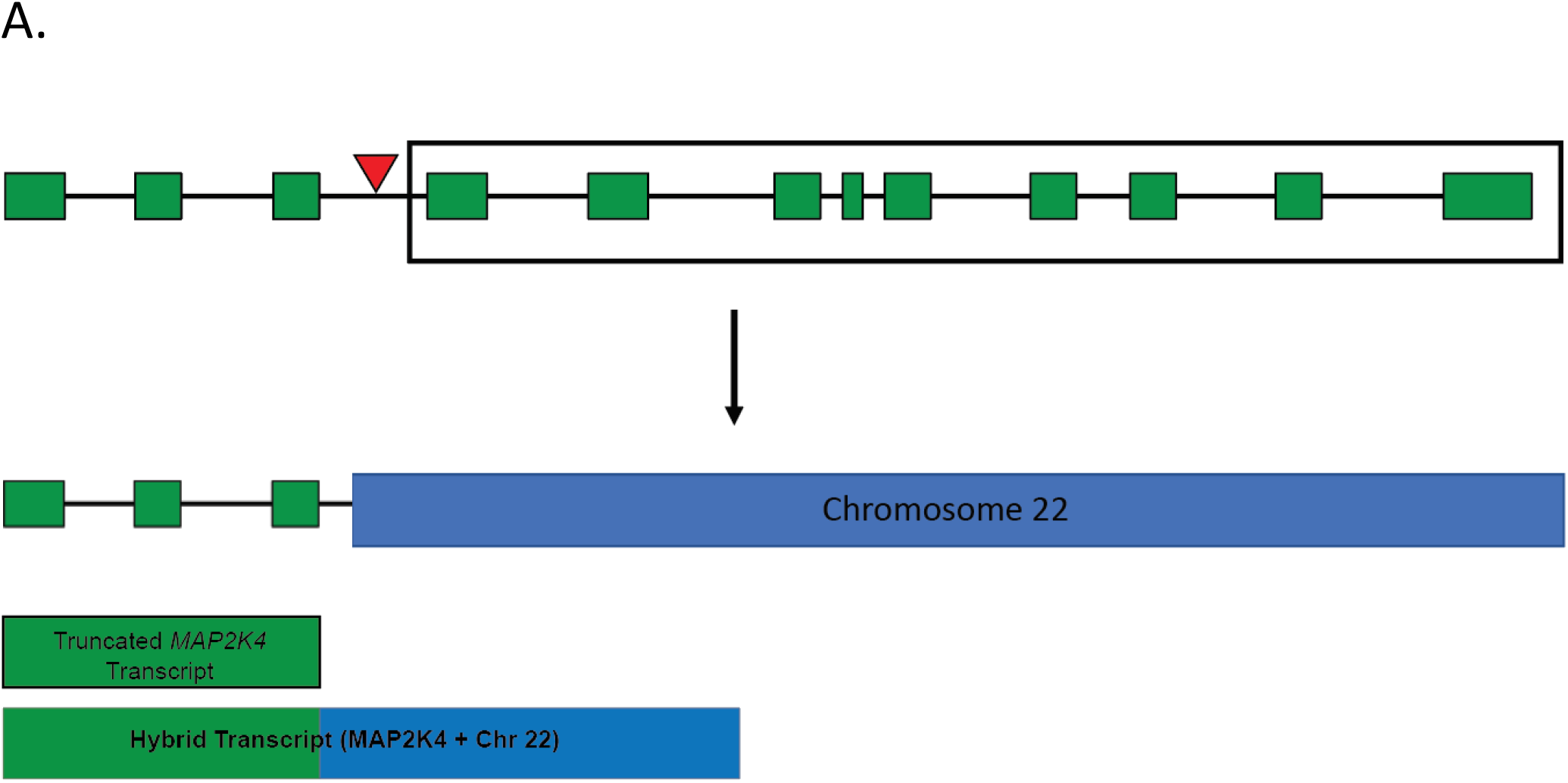
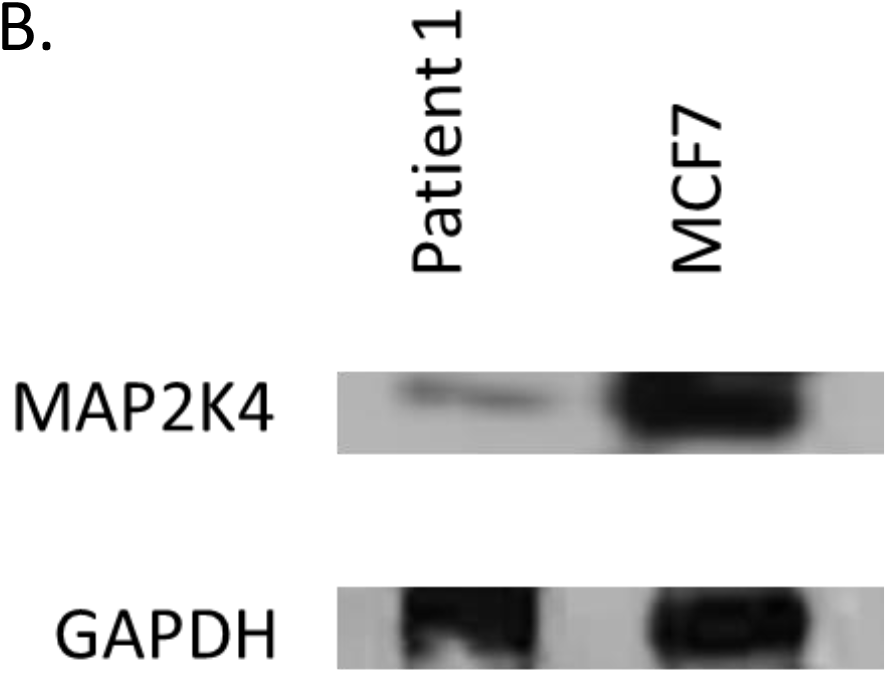
Translocation within *MAP2K4* gene decreases protein production in Patient 1. A) Schematic of the primary transcript of *MAP2K4*. The exons of the transcript are shown as green boxes and the red triangle above *MAP2K4* depicts the approximate location of the translocation. The translocation results in a truncation of *MAP2K4*, leaving only the first three exons remaining on Chromosome 17. The portion of *MAP2K4* shown in the box is translocated to Chromosome 22. The translocation is depicted below the arrow. Truncated and hybrid transcripts are shown below the translocation. B) Western blot of MAP2K4 in control MCF7 cells and ascites-derived cells from Patient 1. The control cells show clear expression of *MAP2K4*, while the cells taken from Patient 1 show decreased production of the protein. GAPDH blotting was performed as a loading control.

Mobile elements can be divided into subfamilies, classified by diagnostic mutations. As different subfamilies may have different expression patterns in cancer, we examined these subfamilies in our patients. Three of the patients in this study had longitudinal bulk RNA-seq data available for further analysis (Supplemental Figure S1). There was no germline RNA-seq data available for these patients, but using SQuIRE we were able to analyze mobile element expression on both subfamily (examining all mobile element transcripts from one mobile element subfamily) and locus-based (specific mobile element loci) levels throughout tumor progression. We saw no subfamily-level change in expression in these three patients for L1, SVA, or *Alu* elements. At the single-locus level, by comparing the four earliest timepoints of Patient 2 to the latter two timepoints (1016 days; nearly 3 years between the fourth timepoint and the fifth), we see significant expression changes (adjusted p-value < 0.05), both increases and decreases, for 337 loci. 125 of these loci are *Alu* elements, 123 are L1s, and 16 are SVA insertions. The remaining loci largely belong to older mobile elements and LTR families. The complete list of mobile element loci that show significantly different expression over time are shown in Supplemental Table S3. Approximately half of the differentially expressed mobile elements overlapped with genes that were also significantly differentially expressed. The location of these mobile elements that overlap with genes is shown in Supplemental Table S4.

The 264 L1s, *Alu* elements, and SVAs that were found to be differentially expressed between the first set of timepoints and second set of timepoints in Patient 2 were intersected with a list of regulatory regions in mammary epithelial tissue. Of these 264 mobile elements, 72 overlapped with a total of 87 regulatory regions. These 87 regulatory regions include a number of CCCTC-binding factor (CTCF) binding sites, open chromatin regions, promoter flanking regions, enhancers, promoters, and transcription factor binding sites. These overlaps are shown in Supplemental Table S5.

## Discussion

Using a trio of mobile element and *de novo* variant detection tools, we identified mobile element insertions and variants in longitudinal WGS data from four breast cancer patients (Supplemental Figure S1). We find that the visual validation step (IGV or other similar tools) reduces the risk of false positive calls and significantly improves the overall accuracy of variant calls (similar to Feusier et al. 2019) (Figure 1). This is particularly important for cancer genomes because most mobile element detection software is not designed for their complexity. This increased complexity makes the visual inspection step necessary, as thousands of potential mobile element insertions suggested by these programs were not unique to the somatic cells, or were the result of mis-mapping in low complexity or mobile element-rich regions. However, the calls from these programs that passed visual inspection spanned both high variant allele fraction and very low variant allele fraction (Figure 2), showing that the insertions or variants did not have to be present at a particularly high frequency to be detected. The SV events identified with this pipeline are largely the result of recombination occurring between mobile elements that are already present in the reference genome (Kolomietz et al. 2002; Gu et al. 2008; Xing et al. 2009) and not a product of somatic retrotransposition (Symer et al. 2002; Gilbert et al. 2005).

Large changes in VAF shown in the current study generally correlate with bottleneck events or large shifts in subclone frequency found previously (Brady et al. 2017). The tumor subclones also change frequency in response to treatment (Brady et al. 2017), which can lead to less stable VAF, a finding shared here when examining mobile element-associated variants over time. The insertions that share similar VAF may be from the same tumor subclone (see the *Alu* insertion and L1 insertion on Chromosome 8 in Patient 2) (Figure 2B), and, after adjusting for tumor purity, those that still show a sharp decline in VAF are likely part of a tumor subclone that showed decreased frequency. The insertions and variants that are present at very high frequency (near 100%) may have occurred early in tumor progression, but these events could also be the result of a bottleneck event or a selective clonal advantage. In cancers with a greater number of these events, mobile elements may be valuable markers for identifying tumor subclones, similar to their role in population genetics and evolutionary studies (Watkins et al. 2003; Witherspoon et al. 2013; Steely et al. 2017; Watkins et al. 2020). Using these markers in conjunction with SNPs may increase the level of resolution for tumor subclones.

From the summary of the insertions and SVs involving mobile elements identified in four patients (Table 1), we show activity for not only L1s, which we expect based on previous studies (Lee et al. 2012; Helman et al. 2014; Tubio et al. 2014; Doucet-O’Hare et al. 2016; Li et al. 2020; Rodriguez-Martin et al. 2020), but also activity from *Alu* and SVA elements. Most previous studies have largely focused on L1s and the structural variants that they mediate (Helman et al. 2014; Tubio et al. 2014), though others have identified a small number of *Alu* insertions (Lee et al. 2012). SVA has been found in a previous study (Helman et al 2014), and recent work also supports post-zygotic insertions in normal tissue for this mobile element family (Feusier et al. 2019). The two SVA insertions identified here are present at low frequency and may have been missed by different filtering methods. We also see variation among patients, with most individuals showing multiple insertions or variants but one showing only a single event that appears to involve an interaction between existing *Alu* elements.

The majority of the variants we identified appear to occur early in our sampling timeframe (Table 1, Figure 2) and likely occur early in tumor progression or development. This suggests that mobile elements may be more active early in cancer and could play a role in tumorigenesis. This supports previous work done in metastatic tumors from multiple patients showing that most insertions that occurred in the primary tumor were reflected in metastases (Ewing et al. 2015). However, ascertainment bias may allow for the increased possibility of identifying high frequency mobile elements and SVs. Further, given the lower sequencing depth of the germline in Patient 2 and Patient 4, it is possible that some of the identified events in these patients could be mosaic. In addition to the early insertions and SVs we identified, we find two insertions, both SVAs, that insert later in tumor progression. It is unclear if something unique about SVA insertions causes them to be more active later in cancer, or if this is just a result of our small sample size. Future studies with a larger sample size of patients should attempt to identify SVA insertions to determine if the trend of insertions at later timepoints remains.

Our analysis of RNA-seq data in these patients showed very few subfamily-level expression changes for mobile elements through time. While we did not have control RNA-seq data from the germline, we were able to examine the course of expression change through tumor progression. Of the three patients with RNA-seq data, we did not observe any instances of subfamily expression changes for *Alu*, SVA, or L1, and only one patient showed a family-level change in expression. This was linked to an increase in HERV activity, which previous studies have reported for HERV-K in various cancers (Wallace et al. 2014; Ma et al. 2016). We were also able to examine the expression of specific mobile element loci in which there were informative reads. Here, using RNA-seq data for Patient 2, which spanned multiple years, we showed that many loci did change expression levels over time. Approximately half of the mobile element loci that showed statistically significant differential expression overlapped genes that were also significantly differentially expressed. Many of the remaining elements were in non-coding regions, though some were found in regulatory regions. Previous work has shown that changes in mobile element methylation and expression can impact nearby gene expression (Xie et al. 2013; Jang et al. 2019). Some of these expression changes are likely tied to methylation changes due to the high CpG content of mobile elements (Yoder et al. 1997). However, differentiating between transposable element transcripts and other transcripts can be challenging, and the patterns shown here may be more indicative of general gene expression change than changes in mobile element expression. We were only able to demonstrate expression changes in Patient 2, who had the most RNA-seq timepoints over the longest course of time. Future studies should examine locus-level expression over time, with germline control comparisons to determine how these expression levels compare with normal cells over longer sampling timepoints.

The early SV events identified in Patient 1 both appear to be associated with *Alu* elements and to affect the coding sequence. The affected genes (*MAP2K4* and *YTHDF2*) have both been implicated in cancer development (Su et al. 2002; Ahn et al. 2011; Koboldt et al. 2012; Xue et al. 2018; Zhong et al. 2019). The deletion we uncovered in *YTHDF2* removes the final exon that would be present in the primary transcript. Though there have been reports of this gene acting as a tumor suppressor, it is not listed in the Catalogue of Somatic Mutations in Cancer (COSMIC) (Tate et al. 2018), and further work likely needs to be completed to validate its role in cancer. The translocation that we identified in Patient 1 interrupted *MAP2K4*, a gene with far more supporting evidence that suggests it plays a role in catalyzing tumor development or metastasis (Teng et al. 1997; Ahn et al. 2011; Pavese et al. 2014). *MAP2K4* is expressed in mammary tissue (median 18.05 TPM; the GTEx Portal on 08/28/2020), and missense and nonsense mutations have been identified at low frequency as part of The Cancer Genome Atlas (TCGA Research Network:https://www.cancer.gov/tcga). The translocation breakpoint on Chromosome 17 occurs after the third exon of *MAP2K4* (Figure 4A), with the split reads in this region showing evidence of an *Alu* element. The breakpoint on Chromosome 22 disrupts a reference *Alu* element, and it is likely that this translocation was associated with an *Alu* interaction. There have also been previous examples of translocations and recombination events associated with mobile elements in cancer (Onno et al. 1992; reviewed in Elliott et al. 2005). *MAP2K4* is listed in COSMIC as potentially having a role in both tumor suppression and as an oncogene, depending on its expression level.

We find decreased copy number along multiple regions of Chromosome 17, including the region that contains *MAP2K4* (Figure 3). Our findings support a two-hit model, where a copy of *MAP2K4* was lost, and the remaining allele was disrupted by the *Alu-*associated translocation described here. With only a single copy of this gene remaining, the translocation renders *MAP2K4* inactive, leading to the decrease in production of the protein shown in the Western blot (Figure 4B). Further analysis of the RNA-seq data for this patient showed that there were no complete transcripts for *MAP2K4. MAP2K4* is found in the *JNK* pathway, which is responsible for numerous cellular functions (Johnson and Lapadat 2002). Previous work in breast cancer tissue has identified multiple mutations along this pathway (including *MAP2K4* and *MAP2K7*) (Koboldt et al. 2012) and suggests that this mutation could act as a driver by altering function of the *JNK* pathway. Brady *et al*. first identified this structural variant in *MAP2K4* in this patient, but by analyzing mobile elements in these patients we have been able to better understand the cause of the mutation.

Overall, we find that most mobile element insertions and the structural variation events (between reference mobile elements) they mediate appear to occur early in tumor development, and most of these early events appear to be passenger mutations. In addition to these passenger mutations, we find SV events involving mobile elements that disrupt the coding sequence of known (*MAP2K4)* and suspected (*YTHDF2)* driver genes in breast cancer. We identified a number of L1, *Alu*, and SVA insertions occurring during tumor progression. As most studies attempt to identify only L1 insertions and structural variants involving L1s, we may be underestimating the impact that mobile elements have on mediating driver mutations in cancer. However, our sample size is small and may not be representative of the activity of these mobile elements in a larger sample size. Future studies should examine other types of cancer for the patterns and impact of mobile element insertions to determine which cancers have an increase in *Alu* and SVA activity, as others have done with L1 insertions. Improving our understanding of mobile element insertions and structural variation events in cancer could enhance our ability to identify tumor subclones and our understanding of the mutational landscape in cancer.

## Methods

### Sequencing data

Information regarding the acquisition of patient sequencing data, as well as quality control and alignment information, can be found in Brady *et al*. 2017. Blood-derived DNA was sequenced and used as the germline DNA sample. Tumor samples were sequenced from ascites and pleural fluid surrounding the breast tumor. DNA sequencing data had previous been aligned to hg19. Data from the Brady *et al*. 2017 publication are available with controlled access on the European Genome-phenome Archive (EGA) under accession EGAS00001002436. Coverage for each WGS timepoint was calculated using covstats from the goleft package (https://github.com/brentp/goleft).

### Mobile element insertions and variants identification

We used the Mobile Element Locator Tool (MELT) (Version 2.1.5) (Gardner et al. 2017), RUFUS (https://github.com/jandrewrfarrell/RUFUS), where possible, and TranSurVeyor (Rajaby and Sung 2018) for the identification of mobile elements and related structural variants in our longitudinal breast cancer samples. Each of these three tools incorporates different methods for identifying variants. Because these tools use different methods, we required only a single tool to identify a potential insertion. Insertions and variants identified by these programs were filtered to include only those that appeared absent in the germline sequencing data. MELT identified 8,825 variants, while TranSurVeyor identified a total of 69,394 variants in these four patients. The total number of variants identified with RUFUS was not calculated because it is designed to look for many types of variants, not only mobile elements. Visual inspection was performed to ensure that the sequences matched a known mobile element sequence using the Integrated Genome Viewer (IGV, Version 2.4.13) (Robinson et al. 2011). Through this visual inspection step, > 70,000 IGV images (as there were multiple individuals included in the MELT analysis) were examined from MELT and TranSurveyor. This step helped to prevent the inclusion of reads that had simply mis-mapped to an incorrect region of the genome. Those insertions that were already present in the germline DNA sample, or those loci that did not show any signs of mobile element activity (low complexity repeats or poorly sequenced regions), were discarded. Reads that appeared to have discordant and/or split reads that mapped to a mobile element were included for further validation. To ensure that the potential mobile element insertions or mobile element-associated structural variants were, in fact, mobile element sequences, we used both BLAT (Kent 2002) and RepeatMasker (Smit 2013-2015). Where possible, we used classic hallmarks of retrotransposition events (target site duplications and poly(A) tails) to ensure that we had identified an insertion. We validated candidate structural variation events that appeared to involve mobile elements with Lumpy (Version 0.2.13) (Layer et al. 2014), and we used IGV validation to identify signs of mobile element involvement (breakpoints in or near existing mobile elements, and small regions of microhomology). While the limit of detection of this pipeline is largely determined by the mobile element detection tools, we were able to identify mobile elements and SVs with VAFs of ∼5%. BEDTools (Quinlan and Hall 2010) intersect was used to determine if the identified variants overlapped with ultraconserved non-coding regions or regulatory elements. Ultraconserved elements and regulatory blocks locations were obtained from UCNEbase (https://ccg.epfl.ch/UCNEbase/) (Dimitrieva and Bucher 2013). The UCSC Genome Browser (ENCODE Regulation track) was manually checked to ensure no other regulatory elements overlapped the insertion site.

### Variant allele fraction

Variant allele fraction was calculated by counting the number of reads showing evidence of the insertion or variant in IGV and dividing this by the total number of reads at the breakpoint of the insertion or variant. The variant allele fraction was then adjusted by multiplying the total number of reads at the breakpoint by the tumor purity (or cancer cell fraction) value. Following this, the number of reads containing the variant were divided by the adjusted estimate of the total number of reads that were derived from tumor cells. The tumor purity estimates were calculated using FACETS (Shen and Seshan 2016), which utilizes SNPs in germline and tumor samples, as well as copy number changes, to provide an estimate of the proportion of tumor cells in the sample. FACETS was also used to determine the location of copy number changes throughout the genome, and particularly at the loci at which we identified variants.

### RNA-seq analysis and mobile element expression

Mobile element expression was measured on a subfamily-specific level and a locus specific level using SQuIRE (Version 0.9.9.9a-beta) (Yang et al. 2019). We compared the first timepoint for each patient to later timepoints to understand how expression patterns changed throughout cancer progression for both subfamily-level and locus-level expression. Additionally, where possible, we compared multiple early sampling timepoints with later sampling timepoints to determine if there was a change during cancer progression. SQuIRE was run according to the documentation provided. Briefly, RNA-seq data were aligned to hg38 using STAR (Version 2.5.3 a) (Dobin et al. 2013); counts of gene expression and mobile element expression were then generated to quantify expression. DESeq2 (Version 1.16.1) (Love et al. 2014) was then used to generate calls for differentially expressed subfamilies and loci. For the locus-specific expression changes at mobile element loci, we intersected these loci with mammary epithelial tissue regulatory regions from Ensembl (Aken et al. 2016) using BEDTools.

Trinity (Version 2.8.5) (Grabherr et al. 2011) was run on bulk RNA-seq data from the first timepoint of Patient 1 to create a library of *de novo* transcripts. The resulting transcripts were aligned to the transcript of *MAP2K4* using BLAT. Transcripts that aligned to *MAP2K4* were further examined to determine which portions of the *MAP2K4* transcript were being produced. Transcripts that aligned to multiple regions of the genome with >95% accuracy were not included.

Salmon (Version 1.4.0) (Patro et al. 2017) was run on RNA-seq timepoints from Patient 1 and Patient 2. The data were converted to gene-level annotations, and the transcripts per million (TPM) for *MAP2K4* were counted. The TPM values for each timepoint for Patient 1 were averaged, and the process was repeated for three timepoints in Patient 2.

### DNA extraction and PCR validation

For validation of our detection methods, PCR amplification was run on a potential L1 and a potential *Alu* insertion in Patient 2, the only patient for whom we had blood-derived germline DNA and were able to extract DNA from cancer cells (ascites). DNA extraction was performed using Qiagen DNeasy Blood and Tissue Kit (50) (Cat No./ID: 69504). PCR amplifications of 25 ng of germline DNA and 25 ng of tumor DNA were performed in 25-µL reactions using Phusion Hot Start Flex DNA polymerase. Initial denaturation was performed for 30 seconds at 98°C, with 40 cycles of denaturation for 10 seconds at 98°C, the optimal annealing temperature of 60°C for 30 seconds, followed by a 2-minute extension at 72°C, and a final extension for 5 minutes at 72°C. The reaction was performed with a negative control (water), the tumor DNA, and the matched germline DNA. Amplicons were run on a 2% agarose gel with ethidium bromide for approximately 90 minutes at 100 V. The gel was imaged using a Fotodyne Analyst Investigator Eclipse imager. The primer sets used for these reactions are listed in Supplemental Table S6, and the corresponding gel images are shown in Supplemental Figures S4 and S5.

### Immunoprecipitation and Immunoblotting

Patient ascites samples were grown in Human Breast Epithelial Cell Culture Complete Media (Celprogen M36056-01S); MCF7 whole cell lysate was purchased (Novus). Cells were lysed with ThermoFisher Co-IP lysis buffer. MKK4 (1:100 CST) and GAPDH (1:300 ProteinTech) antibody were incubated overnight at 4C with 1000µg of protein lysate. Immunoprecipitation was performed according to Pierce™ Classic Magnetic IP/Co-IP Kit protocol. Twenty-eight ug of immunoprecipitated protein was loaded per well of NuPAGE 10% Bis-Tris protein gels. After transfer to PVDF membrane, membranes were blocked with 5% milk:PBST, then incubated with primary antibody (MKK4 1:500, GAPDH 1:000 ThermoFisher) overnight at 4°C. Blots were washed with PBST and incubated with goat anti-rabbit conjugated to HRP secondary antibody for 60 minutes at room temperature. Blots were then incubated in chemiluminescent substrate enhancer (BioRad) and visualized using x-ray film.

## Acknowledgments

This work was supported by the NIH R35GM118335 (to LBJ) and the NIH/NRSA T32HG008962 (to CJS) from the NHGRI. We would like to thank the Biorepository and Molecular Pathology resource at the Huntsman Cancer Institute for their assistance in accessing samples from these patients. We would also like to thank members of both the Jorde and Marth labs for helpful feedback and discussion.

